# Modeling positional effects of regulatory sequences with spline transformations increases prediction accuracy of deep neural networks

**DOI:** 10.1101/165183

**Authors:** Žiga Avsec, Mohammadamin Barekatain, Jun Cheng, Julien Gagneur

## Abstract

**Motivation:** Regulatory sequences are not solely defined by their nucleic acid sequence but also by their relative distances to genomic landmarks such as transcription start site, exon boundaries, or polyadenylation site. Deep learning has become the approach of choice for modeling regulatory sequences because of its strength to learn complex sequence features. However, modeling relative distances to genomic landmarks in deep neural networks has not been addressed.

**Results:** Here we developed spline transformation, a neural network module based on splines to flexibly and robustly model distances. Modeling distances to various genomic landmarks with spline transformations significantly increased state-of-the-art prediction accuracy of *in vivo* RNA-binding protein binding sites for 114 out of 123 proteins. We also developed a deep neural network for human splice branchpoint based on spline transformations that outperformed the current best, already distance-based, machine learning model. Compared to piecewise linear transformation, as obtained by composition of rectified linear units, spline transformation yields higher prediction accuracy as well as faster and more robust training. As spline transformation can be applied to further quantities beyond distances, such as methylation or conservation, we foresee it as a versatile component in the genomics deep learning toolbox.

**Availability:** Spline transformation is implemented as a Keras layer in the CONCISE python package: <u>https://github.com/gagneurlab/concise</u>. Analysis code is available at goo.gl/3yMY5w.

## 1 Introduction

In recent years, deep learning has proven to be powerful for modeling gene regulatory sequences. Improved predictive accuracies have been obtained for a wide variety of applications spanning the modeling of sequences affecting chromatin states (Zhou and Troyanskaya, 2015; Kelley *et al.*, 2016), transcription factor binding (Alipanahi *et al.*, 2015), DNA methylation (Angermueller *et al.*, 2017), and RNA splicing (Leung *et al.*, 2014; Xiong *et al.*, 2015), among others. Using multiple layers of non-linear transformations, deep learning models learn abstract representations of the raw data and thereby reduce the need for handcrafted features. Moreover, the deep learning community, which extends much beyond the field of genomics and includes major web companies, is actively developing excellent software frameworks that allow rapid model development, model exchange, and scale to very large datasets (Collobert *et al.*, 2002; Bastien *et al.*, 2012; Jia *et al.*, 2014; Chollet and Others, 2015; Abadi *et al.*, 2016). Altogether, it is advantageous to leverage these strengths and further develop deep learning modules specific for regulatory genomics.

The distance to defined locations in genes such as transcription start site (TSS), start codon, stop codon, exon junctions, or polyadenylation (polyA) site, which we refer to collectively as genomic landmarks, plays an important role in regulatory mechanisms. Genomic landmarks are often bound by regulatory factors. For instance, RNA 5’ends are bound by capping factors, exon junctions by the exon-junction complex, and the polyA-tail by the polyA-binding proteins. These factors provide spatial clues for other factors to be recruited and to interact. Furthermore, distances to genomic landmarks can be important for structural reasons. The relatively well defined distance between the TATA-box and the transcription start site is due to structural constraints in the RNA polymerase complex (Sainsbury *et al.*, 2015). Also, the splice branchpoints are typically localized within 18 to 44 nucleotides of the acceptor site due to specific constraints of the spliceosome (Wahl *et al.*, 2009; Mercer *et al.*, 2015). Therefore splice branchpoints are not only defined by their sequence but also by their distances to the acceptor site. This information can be used to improve prediction of branchpoint location from sequence (Corvelo *et al.*, 2010; Bitton *et al.*, 2014; Signal *et al.*, 2016).

Despite their important role in gene regulation and their successful usage in computational models, distances to genomic landmarks have not been included in deep learning models. Typical sequence-based deep learning models take into account the effects of relative position *within* each sequence (internal position), either by using strided pooling after convolutional layers followed by fully-connected layers or by using weighted sum pooling (Shrikumar *et al.*, 2017). However, modeling effects of internal positions does not cover modeling of positions to genomic landmarks. These are defined externally to the sequence and can lie at very long distances, as in the case of enhancer to promoter distances. Additionally, genomic landmarks might be difficult to discover *de novo* by the model. While categorical genomic region annotation such as promoter, UTR, intron or exon capture relevant spatial information and help improving prediction performances (Stražar *et al.*, 2016; Pan and Shen, 2017), they are still not capturing distances to genomic landmarks quantitatively.

Here we demonstrate the importance of using relative distances to genomic landmarks as features in sequence-based deep learning models. Technically, we achieve this by introducing spline transformation, a neural network module to efficiently integrate scalar features such as distances into neural networks. Spline transformation is based on smooth penalized splines (P-splines; Eilers and Marx (1996)) and can be applied both in the context of fully-connected layers as well as convolutional layers. We show that deep neural networks modeling effects of distances to genomic landmarks outperforms state-of-the art models on two important tasks. First, we obtain consistent improvements for predicting UV crosslinking and immunoprecipitation (CLIP) peaks across two datasets: a large enhanced CLIP (eCLIP) ENCODE dataset containing 112 RBPs (Van Nostrand *et al.*, 2016) and a well-studied CLIP benchmark dataset (Stražar *et al.*, 2016; Pan and Shen, 2017) containing 19 RBPs from 31 experiments. Second, we obtain the best model for predicting splice site branchpoint (Mercer *et al.*, 2015). Furthermore, we show that across our applications, spline transformation leads to better predictive performance, trains faster and is more robust to initialization than piecewise linear transformations, an alternative class of functions based on the popular rectified linear units.

## 2 Methods

### 2.1 Spline transformation

#### 2.1.1 Definition

We considered input data that not only consist of one-hot-encoded sequence vectors but also of scalar vectors. One typical and simple case is where each input consists of a nucleic acid sequence and a scalar vector of the same length containing the distance of every nucleotide to a genomic landmark of interest (Figure 1). Another case is to have a single value per input sequence, for instance encoding the distance of the sequence midpoint to a genomic landmark. A single value per sequence may be appropriate when positional effects vary over much longer scales than the length of the sequence.

**Figure 1.**
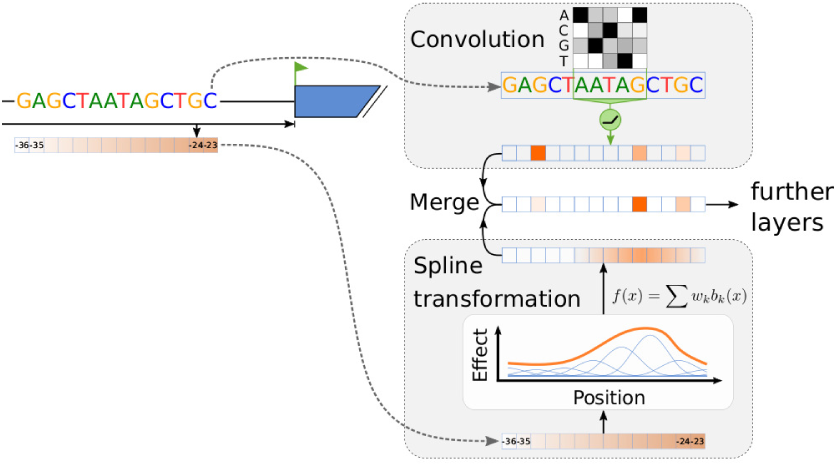
Illustrative model architecture using spline transformation. In addition to DNA sequence, relative distance to various genomic landmarks are used as features. Spline transformation learns a smooth transformation of the raw distances. Transformed distances are then merged with sequence-based activations of convolutional layers.

The positional effects are modeled with a smooth transformation function. We used P-splines or penalized splines (Eilers and Marx, 1996). Spline transformation *f*_*S*_ is defined as

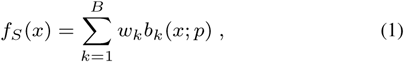

where *b*_*k*_ is the *k*th B-spline basis function of degree *p ∈* ℕ (De Boor, 1978) (Figure 1) and *x* is a multi-dimensional array of positions. In all the applications presented here we used cubic splines, i.e. *p* = 3. Spline bases are non-negative functions with finite support. Knots of the spline basis functions {*b*_1_, *…, b*_*D*_} are placed equidistantly on the range of input values *x*, such that the following relation holds:

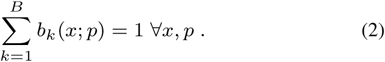

The only trainable parameters in spline transformation are *w*_1_, *…, w*_*B*_.

To favor smooth functions, a smoothness regularization is added to the global loss function:

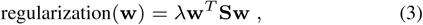

where **S** is a symmetric positive matrix effectively encoding the squared second order differences of the coefficients **w**, which approximate the square of second order derivatives (Eilers and Marx, 1996). The advantage of this approach is that one can have finely spaced bases and use the regularization parameter *λ* to set the amount of smoothing.

#### 2.1.2 Integration into neural networks

Spline transformation is applicable at the network input values. In that case, the approximate range of values–a necessary requirement for the knot-placement–is known. How and where the output of spline transformation is merged into the network is highly application specific. In the case of scalar vectors along the sequence, their spline-transformed values are typically merged with the output of the first sequence-based convolutional layer. In the case of single values per sequence, the transformed values are added to the flattened output of the last convolutional layer, right before the fully-connected layers.

#### 2.1.3 Implementation

We implemented spline transformation using Keras (Chollet and Others, 2015), inspired by the MGCV R package (Wood, 2006). The implementation consists of three essential components: i) a pre-processing function encodeSplines which takes as input an array of values *x*, uniformly places *B* spline bases across the range of *x* and computes [*b*_1_(*x*), *…, b*_*B*_(*x*)] for each array element, ii) a Keras layer SplineT effectively performing a weighted sum of the basis functions and iii) a Keras regularizer SplineSmoother penalizing the squared *m*th-order differences of weights along the last dimension (Equation 3, by default second-order). All three components are compatible with 3 or more dimensional input arrays *x*. Altogether this allows flexible usage of spline transformations in Keras models. The code is open source and is part of the CONCISE python package: github.com/gagneurlab/concise.

#### 2.1.4 Alternative to spline transformation: piecewise linear (PL) transformation

As an alternative to spline transformation, we consider a piecewise linear (PL) transformation achieved by stacking two fully-connected layers with rectified linear unit (ReLU) activation (Nair and Hinton, 2010) in-between. Formally:

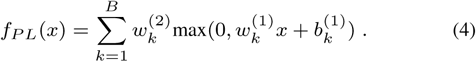

In contrast to spline transformation, the piecewise linear transformation is based on trainable basis functions (max 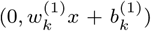) and has hence more parameters. This can be of great advantage when the modeled function is compositional (Montúfar *et al.*, 2014), but can also represent a disadvantage when the modeled function is smooth.

### 2.2 Hyper-parameter tuning with Bayesian optimization

In most of the trained deep neural networks, we employed Bayesian optimization for hyper-parameter tuning using the Tree-structured Parzen Estimator (TPE) algorithm implemented in the hyperopt python package (Bergstra *et al.*, 2013). For each trial, a hyper-parameter configuration is proposed by the Bayesian optimizer. The corresponding model is trained on the training set and evaluated on the validation set. The evaluation metric gets reported back to the optimizer. Model yielding the best performance on the validation set across all trials is selected and evaluated on the test set. This allows for a fair comparison between methods, as all the methods get equal amount of hyper-parameter tuning trials.

### 2.3 eCLIP peak prediction

#### 2.3.1 Data

RNA binding protein (RBP) occupancy peaks measured by eCLIP-seq (Van Nostrand *et al.*, 2016) for human cell lines K562 and HepG2 were obtained from ENCODE version 3 (ENCODE Project Consortium, 2004). There were in total 316 experiments measuring 112 proteins. Genome assembly version GRCh38 and the corresponding GENCODE genome annotation release 25 (Harrow *et al.*, 2012) were used.

For each RBP, a single set of peaks was created by greedily merging the peaks from multiple experiments: two overlapping peaks were merged into one, centered between the peak midpoints. Next, peak midpoints were overlapped with protein-coding genes. Peaks that didn’t map onto any annotated gene (10.1%) were discarded. Each gene-peak pair was considered as a single positive class instance. 94.0% of peaks mapped to a single gene and the average number of mapped genes per peak was 1.064. For each RBP, the negative set was generated by uniformly sampling within each gene 4 times as many locations as true binding sites in that gene. All peaks (both negative and positive) were resized to the width of 101 nt anchored at the peak center. Negative peaks that overlapped positive peaks of the same RBP were discarded.

Finally the sequence underneath the peak was extracted, reverse-complemented for peaks from the negative strand and one-hot encoded. Relative distances from the peak center to the following 8 nearest genomic landmarks on the same strand were extracted: gene TSS, transcript TSS, start codon, exon-intron boundary, intron-exon boundary, stop codon, transcript polyA site and gene polyA site. These features were further transformed with *f*_*pos*_(*x*) = sign(*x*) log_10_(1 + |*x|*) and min-max scaled to fit the [0, 1] range. Data points from chromosomes 1, 3 were used for model validation (17*%*) and hyper-parameter tuning, points from chromosomes 2, 4, 6, 8, 10 (20*%)* for final performance assessment and the rest for model training.

#### 2.3.2 Models

As a baseline model we considered an elastic-net model with *α* = 0.5 (glmnet package, Friedman *et al.* (2010)) based on k-mer counts (*k ∈* 6, 7) and positional features transformed by 10 B-spline basis functions. Smoothness regularization of B-spline features was not used. Optimal number *k* and the regularization strength were determined by 10-fold cross-validation. Models with and without the positional features were compared.

Next, we used a deep neural network (DNN) based on two different data modalities: i) 101 nt one-hot encoded RNA sequence beneath the peak and ii) signed log-transformed relative distances to genomic landmarks. DNN sequence module consisted of two 1D convolutional layers (16 filters each, kernel sizes 11 and 1, ReLU activation after each), followed by max-pooling (pooling size of 4). The positional features were either not used (DNN), or were modeled using spline transformation (DNN w/ dist). Activation arrays of the convolutional layers (RNA sequence) and spline transformation (positional features) were concatenated and followed by two fully-connected layers: a hidden fully-connected layer (100 units and ReLU activation) and a final fully-connected layer with sigmoid activation. Batch normalization was used after every layer. The model was optimized using ADAM (Kingma and Ba, 2014). Bayesian optimization (Section 2.2) was used to determine the optimal set of hyper-parameters for each RBP individually from 20 parameter trials, yielding the best auPR on the validation set.

### 2.4 iDeep CLIP benchmark

#### 2.4.1 Data

To compare our approach with the RBP binding site prediction model iDeep (Pan and Shen, 2017), we used the same CLIP dataset, pre-processing code and model code as Pan and Shen (2017), both provided by the authors at <u>https://github.com/xypan1232/iDeep</u>. The CLIP dataset contains 31 CLIP experiments measuring 19 different RNA binding proteins (RBPs) and was originally generated by Stražar *et al.* (2016) (available at https://github.com/mstrazar/iONMF/tree/master/datasets). Unlike eCLIP (Section 2.3.1), the peaks for each RBP from different experiments were not merged. Respectively, the results are always reported for each experiment individually rather than each RBP. We extended the existing set of features with relative distances to 8 nearest genomic landmarks (gene TSS, transcript TSS, start codon, exon-intron boundary, intron-exon boundary, stop codon, transcript polyA site and gene polyA site), following the same procedure as for the eCLIP data (Section 2.3.1). In contrast to the eCLIP data processing, we used hg19-based GENCODE annotation v24.

#### 2.4.2 Models

As the baseline model we used the provided iDeep model1 with one minor modification: we replaced the softmax activation of the last layer with a sigmoid activation function (softmax is unnecessary for a binary classification task). The iDeep model is based on 5 different data modalities: i) Region type, ii) Clip-cobinding, iii) Structure, iv) Motif and v) Sequence. The additional data modality introduced here —relative distance to 8 genomic landmarks (Section 2.4.1)—was modeled with spline transformation using *B* = 32 basis functions and 6 output units for each features, followed by a fully-connected layer with 64 output units. This module was integrated into the iDeep model by concatenating the activations to the last hidden layer. Spline transformation was used without smoothness regularization, because we restricted ourselves the same set of hyper-parameters as the iDeep model and have not done any hyper-parameter tuning. All models were optimized using RMSprop (Tieleman and Hinton, 2012), same as the original iDeep.

### 2.5 Branchpoint prediction

#### 2.5.1 Data

Branchpoint prediction is a binary classification task to predict measured high-confidence branchpoints in introns, 18-44 nucleotides upstream of the 5’ intron-exon boundary. The same dataset and pre-processing procedure was used as described in Signal *et al.* (2016). Briefly, high-confidence annotated branchpoints from Mercer *et al.* (2015) were used to generate the positive set. Negative set comprises of positions not annotated as high-or low-confidence branchpoints in Mercer *et al.* (2015). This yields in total 52,800 positive and 933,739 negative examples. Signal *et al.* (2016) designed and used the following features in the classification model (Figure 3a): 11 nucleotide sequence window around the position encoded as dummy variables, distances to the first 5 canonical AG dinucleotides downstream, distance to the poly-pyrimidine tract (PPT) and its length, distance to the associated 3’ exon and distance to the nearest 5’ exon located on the same strand. GENCODE v12 (Harrow *et al.*, 2012) was used for genome annotation. Using the code provided by Signal *et al.* (2016)2, we were able to reproduce the results of Signal *et al.* (2016). The only major change to the pipeline was to *a priori* set aside points from chromosomes 4, 5, 6, 7, 8, and X (21% of all the data) as a test set. The test set was only used to test the predictive performance of our models and not to tune the hyper-parameters as done in Signal *et al.* (2016). Exact code changes can be tracked in our forked repository3.

#### 2.5.2 Models

All the models use the same set of features as Signal *et al.* (2016).

**branchpointer** - branchpoint prediction model developed by Signal *et al.* (2016). It is a combination of two stacked models: SVM with “rbfdot” kernel and a gradient-boosted decision trees model, both from the caret R package (Kuhn, 2015).

**glmnet** - Logistic regression with elastic-net regularization using the glmnet R package (Friedman *et al.*, 2010) with parameters alpha=0.5 and regularization strength determined by 5-fold cross-validation on the training dataset.

**NN** - deep neural network developed here (Figure 3b). For computational efficiency, the model predicts the branchpoint class for all 27 positions in an intron simultaneously, while using the same parameters for each position. Specifically, the models takes as input one-hot encoded 37 nt long RNA sequence and 9 position-related features, each as an integer array of length 27. Parameter sharing across 27 positions within an intron is achived with 1d-convolutions using kernel size of 1. The only exception is the first convolutaional layer processing RNA sequence where kernel size of 11 is used. That way, the set of features for predicting the branchpoint class at a single position is exactly the same as for branchpointer and positions are completely independent of each other.

The 9 positional features were transformed either with: i) spline transformation (ST) or ii) piecewise linear transformation (PLT). Moreover, two levels of model complexity were compared: ‘shallow’ and ‘deep’. They differ in the number of convolutional filters, number of hidden layers (Figure 3b) and also in the weight initialization for the first sequence-based convolutional layer: ‘shallow’ models were initialized with the position-specific scoring matrix (PSSM) of the high-confidence branchpoints derived from the training set and ‘deep’ models were initialized with the (random) glorot-uniform initialization. In total, 4 different model architectures were used. Hyper-parameters were tuned for each of the four NN classes individually using Bayesian optimization (Section 2.2).

#### 2.5.3 Position-weight matrix analysis

Weights of the convolutional filter *w*_*ij*_ were converted to a position-weight matrix (PWM) by

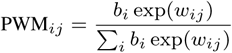

where *i ∈ {A, C, G, T}* is the nucleotide identity and *b*_*i*_ is the background probability (A: 0.21, C: 0.25, G: 0.20, and T: 0.34 in the branchpoint dataset). Note that we are denoting T also Uracil. Branchpoint-centered PWM was created from 11 nt long sequences centered at the high-confidence branchpoints from Mercer *et al.* (2015).

## 3 Results

### 3.1 Relative distance to genomic landmarks improves *in vivo* RBP binding prediction

We first investigated the benefit of modeling effects of position with respect to genomic landmarks for the task of predicting *in vivo* binding sites of RNA-binding proteins (RBPs). We used a large and consistently generated dataset of eCLIP data for 112 RBPs from the ENCODE project (Methods, Van Nostrand *et al.* (2016)).

For a representative detailed investigation, we first focused on 6 RBPs with more than 10,000 peaks and exhibiting various peak distributions along genes (Figure 2a). Comparing the relative positions within genes between the binding and non-binding sites (Figure S1), we selected 2 RBPs with highest enrichment toward the transcription start site (TSS; DDX3X, NKRF, t-test comparing positions of binding sites versus non-binding sites *P* < 10^−100^), 2 RBPs showing highest enrichment toward the polyA site (UPF1 and PUM2, *P* < 10^−100^) and 2 RBPs showing the least significant positional preference (TARDBP, SUGP2, *P >* 0.5). We next asked what the contribution of using a deep neural network on the one hand and of modeling positional effects on the other hand for the task or predicting eCLIP peaks was. To this end, we fitted four models (Methods): i) an elastic net logistic regression based on k-mer counts from 101 nt sequence around the candidate peak as a non-deep supervised learning algorithm (glmnet), ii) an extension of the latter model that also included relative distance to TSS and polyA site transformed by spline transformation (glmnet w/ dist), iii) a deep neural network based on the 101 nt sequence around the candidate peak (DNN), iv) an extension of the latter model with spline transformation of relative distance to TSS and polyA site (DNN w/ dist).

**Figure 2.**
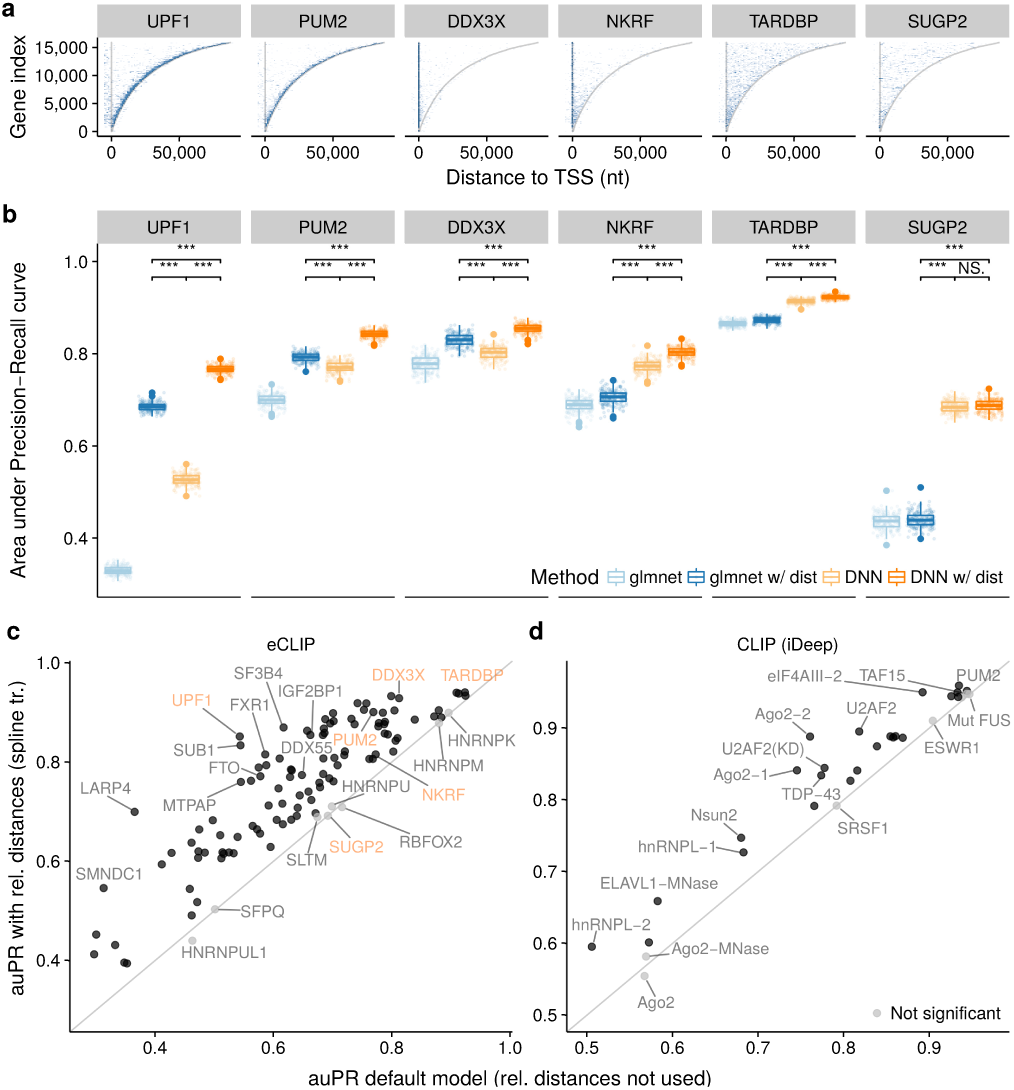
Relative distance to genomic landmarks boosts the *in vivo* prediction of RBP binding sites. **a)** eCLIP peak distribution across all genes. Genes (y-axis) are sorted by their length and aligned at their start site. Color intensity represents the number of peaks per bucket (100 genes x 1000 nt) and saturates at 10 peaks per bucket. Grey lines represent gene transcription-start-site (TSS) and polyadenylation (polyA) site. **b)** Area under Precision-Recall curve (auPR) for predicting in vivo RBP binding sites measured by eCLIP for a subset of RBPs (6/112). Methods labelled by “w/ dist” rely, in addition to RNA sequence, on two positional features: distance to TSS and polyA site. Distribution of the auPR metric (boxplot instead of point-estimate) is obtained by generating 200 boostrap samples of the test-set and computing auPR for each of them. ^***^ denotes *P* < 0.001 (Wilcoxon test). **c,d)** Benefit of adding 8 genomic landmark features with spline transformation to the (c) DNN model for all 112 RBPs measured by eCLIP in ENCODE and (d) iDeep model (Pan and Shen, 2017) for 19 RBPs across 31 CLIP experiments. Black represents statistically significant difference (*P* < 0.0001, Wilcoxon test on 200 bootstrap samples, Bonferroni correction for multiple testing).

**Figure 3.**
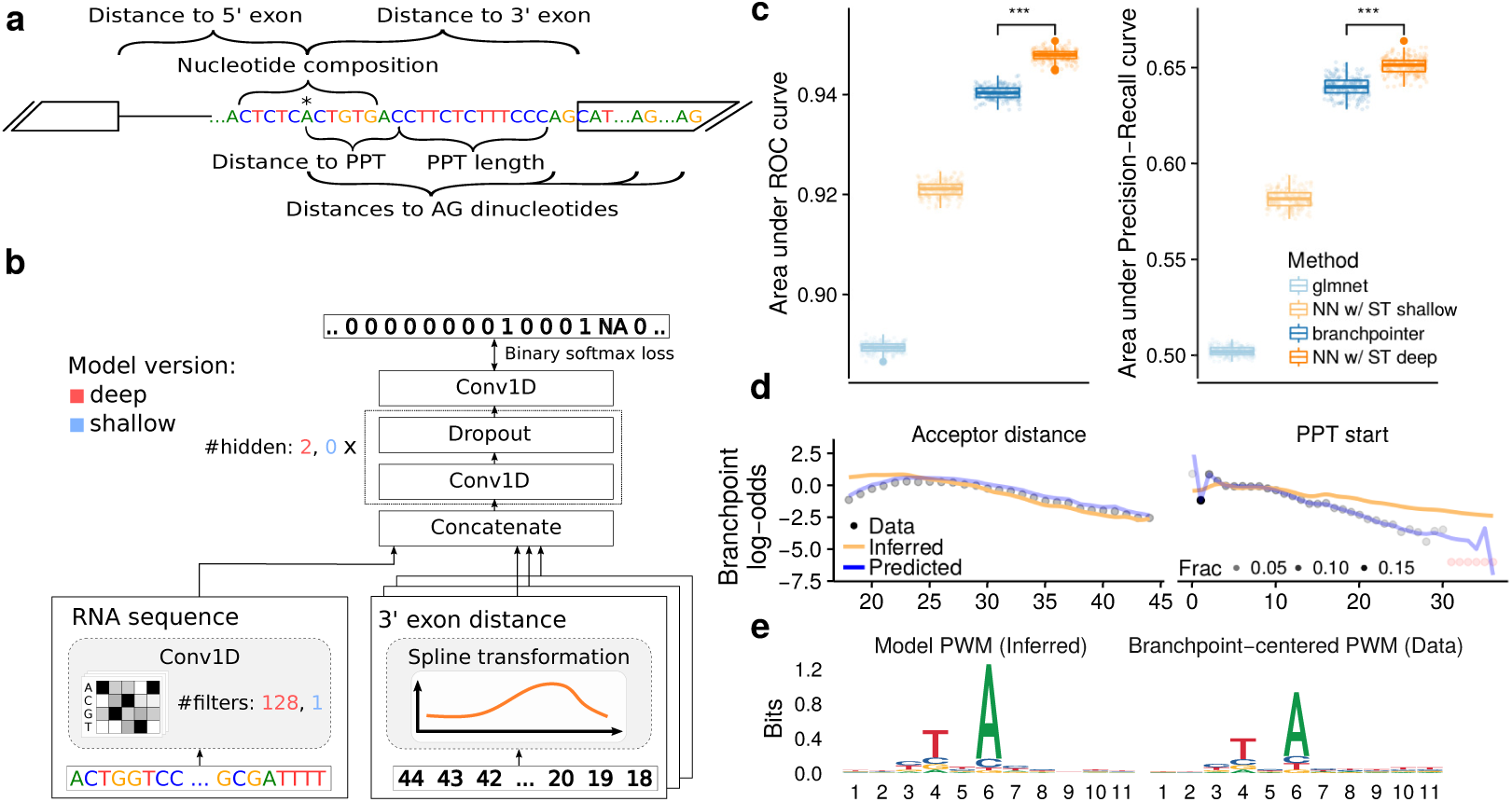
Spline transformation improves branchpoint prediction: **(a)** Features for branchpoint prediction designed by Signal et al. (2016) (adapted from Signal et al. (2016)). PPT stands for poly-pyrimidine tract. **(b)** NN model architectures (deep and shallow) for branchpoint prediction developed here. For each predicted binary class (1=high confidence branchpoint, NA=ignored low-confidence branchpoint, 0=else), the model takes as input 11-nucleotide sequence window and 9 position-related features. **(c)** auROC and auPR bootstrap distribution (n=200) for branchpoint prediction on the test set. Our NN models are compared to the state-of-the-art model branchpointer (Signal et al., 2016) and an elastic-net baseline using the same features. ^***^ denotes *P* < 0.001 (Wilcoxon test). **(d)** Fraction of branchpoints per position for the two most important features in log-odds scale (black dot, outlier shown in red) compared to the shallow NN model fit: inferred spline transformation (orange) and predicted branchpoint log-odds (blue). **(e)** Information content of the shallow NN convolutional filter transformed to the PWM and the branchpoint-centered PWM (Methods).

For each of the 6 RBPs, the deep neural networks yielded a significantly larger area under the precision-recall curve (auPR, a metric between 0 and 1, the larger, the better) compared to their corresponding elastic-net based models (Figure 2b). Moreover, modeling positional effects significantly improved the performance for all four RBPs showing positional preference. In three out of four cases (UPF1, PUM2 and DDX2X), the glmnet model even outperformed the deep neural network model DNN lacking positional features. These results show the importance of modeling positional effects for predicting RBP binding peaks and the power of combining deep neural networks with the modeling of position to genomic landmarks.

Next, we extended our set of positional features in DNN w/ dist to 8 genomic landmarks (nearest gene TSS, transcript TSS, start codon, exonintron boundary, intron-exon boundary, stop codon, transcript polyA site, gene polyA site; Figure S2) and compared it with DNN across all the 112 RBPs. Using relative distances increased the auPR by up to 0.33 (LARP4, from 0.37 to 0.70), on average by 0.11 (from 0.64 to 0.75, *P* < 10^−16^ paired Wilcoxon test, Figure 2c). Altogether, 104 RBPs showed significant auPR increase and none a significant decrease (*P* < 10^−4^, Wilcoxon test, Bonferroni correction for multiple testing).

To further validate our observations from the eCLIP data, we extended the current state-of-the-art model for RBP binding site prediction–iDeep (Pan and Shen, 2017) with the same 8 genomic landmark features. iDeep is a deep neural network trained and evaluated on a CLIP dataset of 19 proteins measured by 31 experiment created by Stražar *et al.* (2016). It does not model distances to genomic landmarks quantitatively. However, it is based on indicator features for 5 gene regions (exon, intron, 5’UTR, 3’UTR, CDS) for each nucleotide in the classified sequence. Since iDeep was already implemented in Keras, extending it with our spline transformation module could be done easily (Methods). When we added the 8 positional features with spline transformation on top of iDeep, the auPR increased by 0.036 (*P* = 4.7 *×* 10^−9^, paired Wilcoxon test) and auROC by 0.021 (*P* = 2.3 *×* 10^−8^). The auPR improved significantly for 25 out of 31 experiments and has not significantly decreased for any experiment (Figure 2d). This shows that the quantitative distance, and not just a binary indicator, are useful predictive features for RNA binding sites. Moreover, this application demonstrates how spline transformation modules can enhance existing deep learning models.

Altogether, these results demonstrate that relative distance to genomic landmarks is an important feature for predicting *in vivo* RBP binding events and show that our spline transformation module provides an effective way to include this information in deep neural networks.

### 3.2 Spline transformation in a deep neural network improves state-of-the-art branchpoint prediction

We then asked whether spline transformation in a deep neural network could improve prediction accuracy for tasks where the effect of the distance to genomic landmarks has already been exploited by non-deep learning methods. To this end, we considered the prediction of splice branchpoint. The first reaction of splicing is the attack of a 2’hydroxyl group of an intron adenosine on the 5’ splice site phosphodiester bond (Ruskin *et al.*, 1985). This intron adenosine is located typically between 18 and 44 nucleotide 5’ to the acceptor site (Mercer *et al.*, 2015). It is named branch point, because it is bound on its 2’hydroxyl group, leading to a lariat form of the spliced-out intron. Mapping branchpoints experimentally has been difficult because of the very short half-life of lariats. Computational predictions of branchpoints have been also difficult because their sequence context is degenerate (Gao *et al.*, 2008).

Current state-of-the art model to predict human branchpoints is *branchpointer* (Signal *et al.*, 2016), an ensemble model of support vector machine (SVM) and gradient boosting machine (GBM) trained on a set of 42,095 mapped high-confidence branchpoints from Mercer *et al.* (2015). In addition to the sequence context, *branchpointer* uses 11 different positional features: distances to the first 5 downstream AG dinucleotides, distance to the poly-pyrimidine tract (PPT) and its length, distance to the associated 3’ and 5’ exon (Figure 3a, Methods).

Using the provided code and some minor modifications (Methods), we were able to reproduce the results of branchpointer and obtained very similar performance metrics as originally reported: area under Receiver operating characteristic curve (auROC) of 0.940 (paper: 0.941) and area under precision-recall curve (auPR) of 0.640 (paper 0.617) (Figure 3c). Training a deep neural network with spline transformation module for positional features (Figure 3b) significantly outperformed *branchpointer* with auROC of 0.949 and auPR of 0.651 (*P* < 2.2 *×* 10^−16^, Wilcoxon test, Figure 3c). This result is consistent with general improved performance of deep neural networks over alternative supervised learning models and shows the strength of spline transformation. It also yields to the most accurate predictor of human branchpoints to date.

### 3.3 Shallow architecture yields an interpretable branchpoint model while still delivering good predictive performance

When model interpretation rather than mere prediction is desired, shallow neural networks are preferred over deep neural networks because their coefficients can be directly interpreted. To investigate such a use case, we trained a shallow version of our neural network (NN w/ ST shallow, Methods) for branch point prediction. As expected, the shallow model is not able to compete with its deeper version or *branchpointer*. Nevertheless, it performs well compared to an elastic-net logistic regression (Figure 3c).

Predicted positional effects in the shallow model (’Predicted’ in Figure 3d, Figure S3) closely resembled the distributions of branchpoint distances to all genomic landmarks (’Data’ in Figure 3d, Figure S3). In addition to the distances, the single convolutional filter in our shallow model captured the expected sequence preference of branchpoints (Figure 3e, Mercer *et al.* (2015)).

Altogether, these analyses of branchpoint prediction demonstrate the versatility of the spline transformation module. The spline transformation module can be used to increase predictive power in conjunction with deep neural network. It can also be employed in shallow and interpretable models.

### 3.4 Spline transformation is more robust to hyper-parameter choices, trains faster, and yields better predictive performance than piecewise linear transformation

The most widely used transformations in deep leaning currently are compositions of linear transformation and rectified linear units, defined as ReLU(x) = *max*(0, *x*). Those compositions lead to piecewise linear transformations (Methods). Although piecewise linear functions can approximate any function, this can be at the cost of much more parameters. Also piecewise linear functions are not smooth.

To inspect the benefit of spline transformation compared to the default modeling choice in deep learning, we replaced the spline transformation module with piecewise linear (PL) transformation (Methods) in all three studied tasks. As for the spline transformation, we used the same number of single output units, equivalent number of hidden units and the same hyper-parameter optimization strategy for each task. The observed predictive performance of spline transformation was consistently better across all 3 tasks (Figure 4a-c): i) for eCLIP the auPR improved on average by 0.042 (*P* < 2.2 *×* 10^−16^, paired Wilcoxon test) and auROC by 0.025 (*P* < 2.2 *×* 10^−16^); ii) for the iDeep CLIP benchmark dataset, the auPR improved on average by 0.011 (*P* = 1.3 *×* 10^−3^) and auROC by 0.004 (*P* = 5.4 *×* 10^−4^); iii) for branchpoint, the auPR improved on average by 0.005 (*P* = 1.0 *×* 10^−12^ and auROC by 0.002 (*P* < 2.2 *×* 10^−16^).

**Figure 4.**
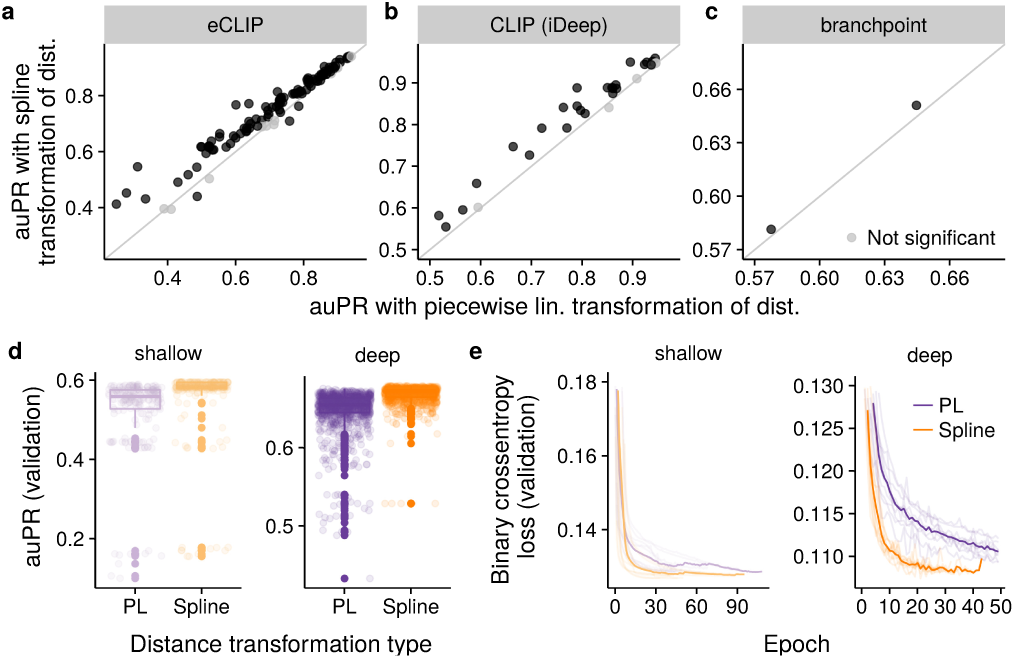
Spline transformation outperforms piecewise-linear transformation in terms of generalization accuracy, hyper-parameter robustness and training efficiency. **(a-c)** Test-accuracy (auPR) comparing spline transformation to piecewise linear transformation for all the tasks presented in the paper (Figure 2c, Figure 2d and Figure 3c). Black represents statistically significant difference (*P* < 0.0001, Wilcoxon test on 200 bootstrap samples, Bonferroni correction for multiple testing). **(d,e)** Training and hyper-parameter tuning metrics for the branchpoint task. **(d)** Validation accuracy (area-under precision-recall curve) of all the hyper-parameter trials. **(e)** Training curves (validation loss per epoch) of 10 best hyper-parameter trials (transparent lines) and their average (solid line).

Focusing on the branchpoint prediction task, we compared the validation accuracies of all hyper-parameter trials between spline transformation and piecewise linear transformation (Figure 4d). Spline transformation had fewer trials with poor performance and globally smaller performance variation. This suggests that spline transformation is more robust to parameter initialization and hyper-parameter choices like the learning-rate. Moreover, we inspected the training curves of the top 10 trials (Figure 4e). While the deep neural network with spline transformation on average trained in 20 epochs, piecewise linear transformation required more than 50 epochs. Altogether these results show that spline transformations generalize better, are more robust and train in fewer steps than piecewise linear transformations in the class of problems we investigated.

## 4 Discussion

Here we have introduced spline transformations, a module for neural networks, and demonstrated that it is an effective way to model relative distance to genomic landmarks. Spline transformations allowed us to improve the state-of-the-art prediction accuracy of splice branchpoint and *in vivo* RBP binding affinity. On the latter task, the use of relative distance to genomic landmarks in a neural network is novel. Moreover, we have shown that spline transformation in a shallow network can uncover the positional effects of cis-regulatory elements.

We provide spline transformation as an open source Keras components, compatible with both Theano (Bastien *et al.*, 2012) and TensorFlow (Abadi *et al.*, 2016). We have shown how to combine it with existing models and improve their performance. Compared to a 2-layer neural network with ReLU activations—piecewise linear transformation, spline transformation offers better prediction accuracy, is more robust to initialization and trains faster. This is not surprising as the relative positional features tend to affect the response variable in a smooth fashion, which is exactly the class of functions spline transformation is able to represent with very few parameters.

In addition to external positions studied here, spline transformation can also be used to model *internal* positions, which are positions within the sequence. In that case, the array index *i* along the spatial dimension of the 1d convolutional layer activation *a*_*ij*_ serves as the relative distance feature. That way, weights in the the recently introduced weighted sum pooling layer (Shrikumar *et al.*, 2017) can be parametrized by spline transformation: *w*_*ij*_ = *f*_*S*_ (*i*). Note that this applies also to the separable fully-connected layer (Alexandari *et al.*, 2017), which can be reformulated as 1d convolution with kernel size of 1 followed by a weighted sum pooling layer. Altogether, using spline transformation for modeling internal position reduces the number of parameters in the network even further.

One limitation of spline transformation is that scale of the input features (e.g. log or linear) remains important and has to be chosen upfront, because spline knots are placed uniformly across the whole range of feature values. We suggest users to perform pre-processing investigations to identify the most appropriate scales for the problem at hand. Moreover, the current implementation of spline transformation is not able to model the interaction between variables directly. Note that while this interaction is still captured by the downstream fully-connected layers, a more appropriate solution might be to use multi-dimensional B-splines. A further research direction is the estimation of confidence bands for the inferred spline transformation function. Confidence bands for spline estimates are available in the context of generalized additive models (Hastie and Tibshirani, 1990). We have recently shown how this can be used to perform differential occupancy analysis of ChIP-seq data (Stricker *et al.*, 2017). Confidence bands would allow deriving statistically supported claims about the positional effects of cis-regulatory elements.

We have demonstrated the power of using spline transformations for modeling effects of distances to genomic landmarks. However, the spline transformation module is more general. It could be used to transform any other relevant scalar. Relevant scalars for cis-regulatory elements include conservation scores, modifications such as methylation rates, experimental measures such as occupancies by factors, or nucleosomes. Hence, we foresee spline transformations as a useful generic tool for modeling of cis-regulatory elements with neural networks.

## Acknowledgements

We thank Basak Eraslan and Roman Kreuzhuber for useful feedback on the manuscript and Bethany Signal for sharing insights regarding her branchpoint analysis.

## Footnotes

1 https://github.com/xypan1232/iDeep

2 https://github.com/betsig/splice_branchpoints

3 https://i12g-gagneurweb.informatik.tu-muenchen.de/gitlab/avsec/splice_branchpoints

---

**Figure S1.**
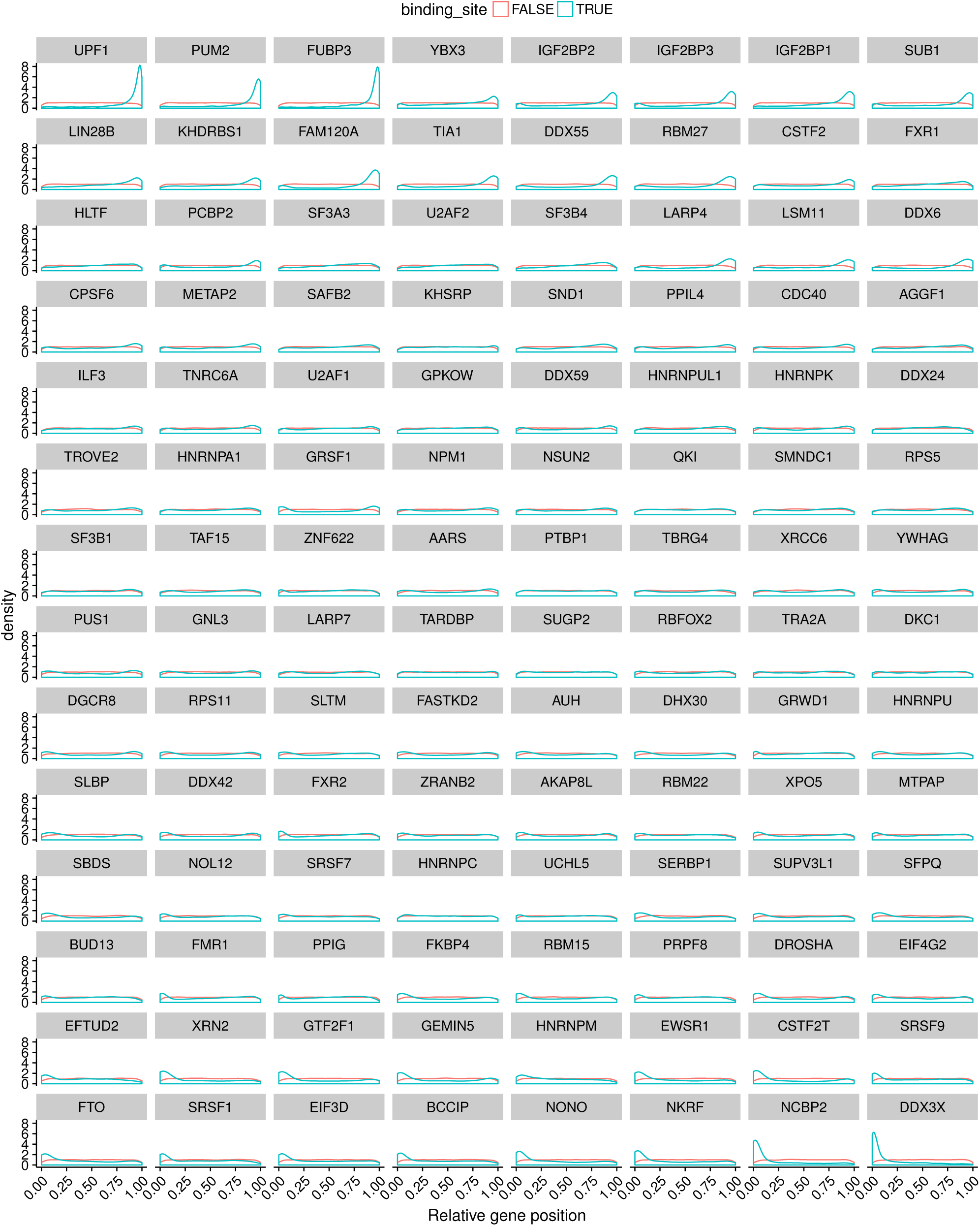
Figure 2 supplement: Relative genes positions for 112 RBPs measured by eCLIP. RBPs are sorted by the t-test statistic comparing relative genes positions of binding to non-binding sites.

**Figure S2.**
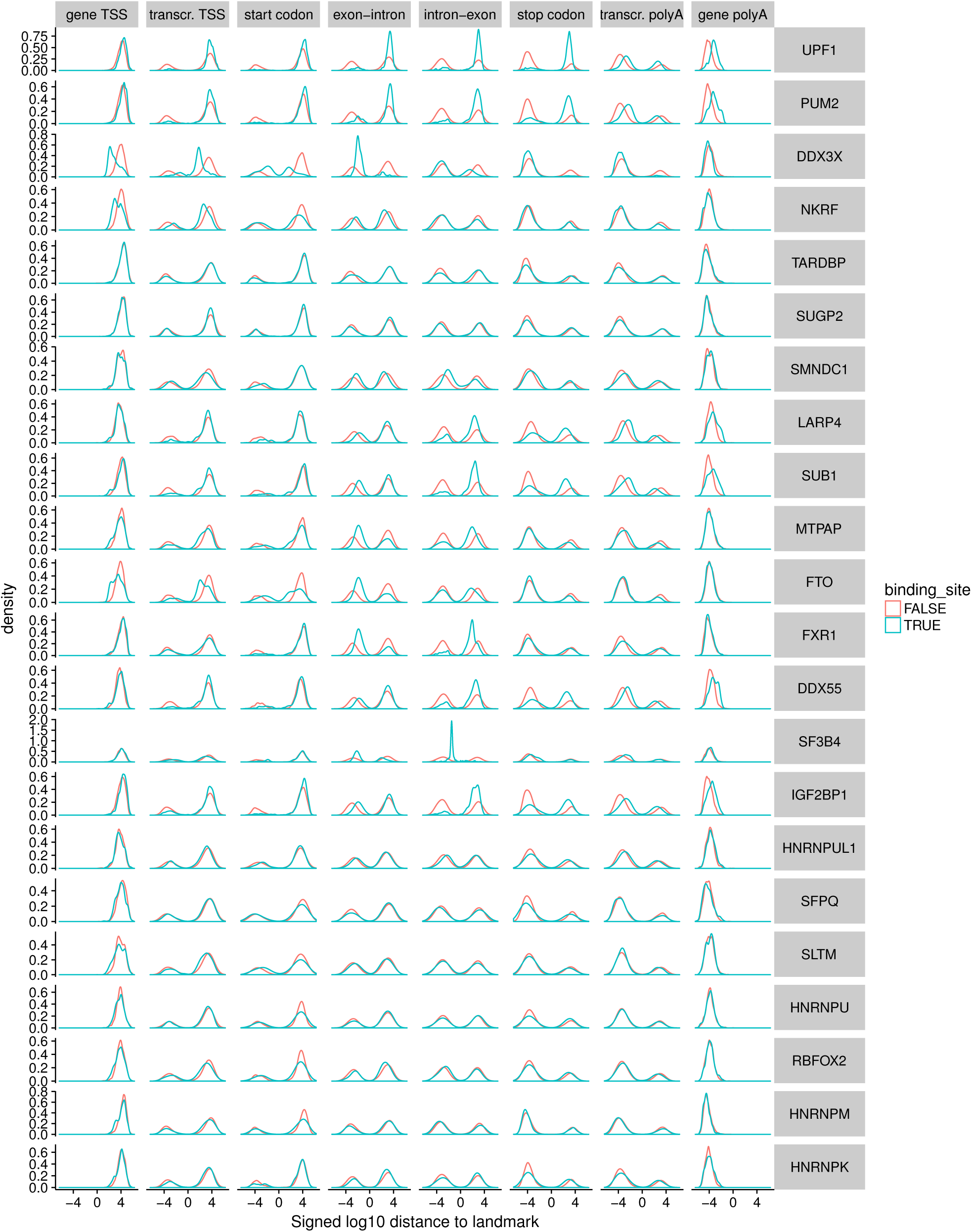
Figure 2 supplement: Relative distance to all 8 considered genomic landmarks for eCLIP peaks. Only RBPs labelled in Figure 2c are shown.

**Figure S3.**
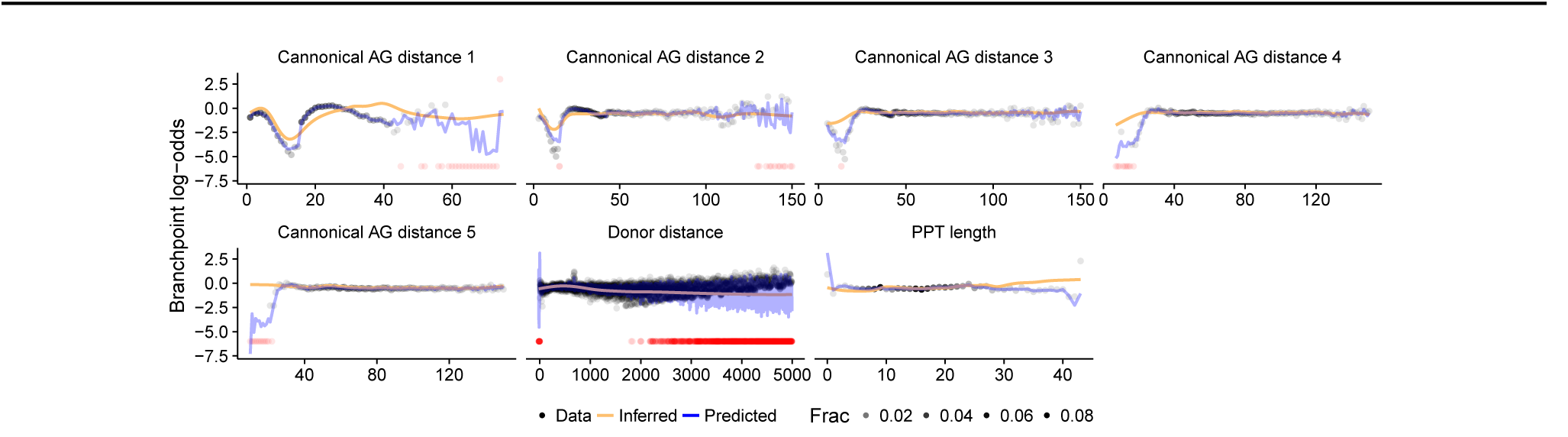
Figure 3 supplement: Fraction of branchpoints per position for the remaining seven features in log-odds scale (black dot, outlier shown in red) compared to the shallow NN model fit: inferred spline transformation (orange) and predicted fraction of branchpoint per position (blue).

